# Incorporating strontium enriched amorphous calcium phosphate granules in collagen/collagen-magnesium-hydroxyapatite osteochondral scaffold improves subchondral bone repair

**DOI:** 10.1101/2023.06.15.545062

**Authors:** Jietao Xu, Jana Vecstaudža, Marinus A. Wesdorp, Margot Labberté, Nicole Kops, Manuela Salerno, Joeri Kok, Marina Simon, Marie-Françoise Harmand, Karin Vancíková, Bert van Rietbergen, Massimiliano Maraglino Misciagna, Laura Dolcini, Giuseppe Filardo, Eric Farrell, Gerjo J.V.M. van Osch, Jānis Ločs, Pieter A.J. Brama

## Abstract

To date, osteochondral defect repair with a collagen/collagen-magnesium-hydroxyapatite (Col/Col-Mg-HAp) scaffold has demonstrated good clinical results. However, subchondral bone repair has been suboptimal, potentially leading to damage to the regenerated overlying neocartilage. This study aimed at improving the bone repair potential of this scaffold by incorporating strontium (Sr) ion enriched amorphous calcium phosphate (Sr-ACP) granules (100-150 µm). Sr concentration of Sr-ACP was determined with ICP-MS at 2.49 ± 0.04 wt.%. Then 30 wt.% ACP or Sr-ACP granules were integrated into the scaffold prototypes. The ACP or Sr-ACP granules were well distributed and embedded in the collagenic matrix demonstrated by micro-CT and scanning electron microscopy/energy dispersive x-ray spectrometry. Good cytocompatibility of ACP/Sr-ACP granules and ACP/Sr-ACP enriched scaffolds was confirmed in *in vitro* cytotoxicity assays. An overall promising early tissue response and good biocompatibility of both ACP and Sr-ACP enriched scaffolds were demonstrated in a subcutaneous mouse model. In a goat osteochondral defect model, significantly more bone observed at 6 months with the treatment of Sr-ACP enriched scaffolds compared to scaffold only in particular in the weight-bearing femoral condyle subchondral bone defect. Overall, the incorporation of osteogenic Sr-ACP granules in Col/Col-Mg-HAp scaffolds showed to be a feasible and promising strategy to improve subchondral bone repair.

## 1. Introduction

Pain and restriction-free movement of joints is possible when the osteochondral unit is well preserved. The native osteochondral unit is composed of two main tissue types: articular cartilage and subchondral bone which are connected via calcified cartilage. A healthy articular cartilage ensures joint lubrication and stress reduction, and the subchondral bone is crucial for underlying mechanical support. These functions can be altered if the complex structure of the osteochondral unit is damaged by traumatic injuries, chronic diseases, and age-related degeneration. Endogenous osteochondral defect repair is limited due to the lack of a vascular/nerve supply in the cartilage and the complex multiphasic structure of the osteochondral unit [1, 2]. Due to its limited self-healing capacity, osteochondral defects may progress into osteoarthritis without effective and timely intervention. To regenerate osteochondral tissues in the lesion site, surgical interventions, such as autologous chondrocyte implantation, osteochondral grafting, and microfracture have been extensively applied. However, the regenerated tissue consists mostly of a mixture of fibrous tissue and fibrocartilage [3, 4], leading to poor resistance to shear forces and deterioration at long term follow-up [5, 6].

To improve osteochondral tissue repair, biomaterial-based scaffolds have shown promising results in regenerating damaged tissues. To mimic the native osteochondral composition and structure, biomaterial-based bilayered scaffolds have been developed and tested [7]. Among these, a scaffold with a superficial collagen-only layer and a deep layer of collagen mixed with hydroxyapatite (HAp) represents a promising substitute [8, 9]. Clinically, this collagen/collagen-magnesium (Mg)-HAp (Col/Col-Mg-HAp) scaffold has demonstrated good stability and clinically relevant improvement in knee function [10–12]. However, subchondral bone repair remained suboptimal in comparison to the cartilage repair capacity of this scaffold in clinical follow-up [12]. An unrepaired subchondral bone may affect the biomechanical properties of the osteochondral unit, which might lead to damage to the regenerated overlying neocartilage and joint pain for the patient. Well-healed subchondral bone is therefore critical to support long-term survival of the overlying neocartilage [13].

We hypothesize that addition of extra calcium phosphate (CaP) to the Col/Col-Mg-HAp scaffold would enhance regeneration of the subchondral bone. That extra CaP could be the well-known hydroxyapatite (HAp, Ca_10_(PO_4_)_6_(OH)_2_) which is a close chemical analogue to the biological apatite present in bone[14]. However, the stoichiometric HAp, in comparison with biological apatite, has low solubility and resorbability [15]. Limitations of HAp could be overcome by using amorphous calcium phosphate (ACP) instead. ACP is a hydrated CaP with an amorphous structure, having possibility of different Ca/P molar ratios (1.2-2.2), and high specific surface area [16]. Presence of an amorphous phase, hydrated structure and high specific surface area of ACP are shared with the biological apatite [17], and it ensures ACP’s bioactivity, solubility, and the excellent adsorption properties[16].

As ACP’s amorphous structure can accommodate other ions besides calcium and phosphate[18], it can be upgraded to include an ion for additional bone regenerative effect. Bioinorganic ions, e.g., strontium (Sr), are cost-effective, easy to use and a local delivery tool[19] having less risks than bone morphogenetic protein (BMP) strategies used for improved regeneration of bone [20]. Previously Sr has been introduced in forms of Sr ranelate drug or as a dopant in the biomaterial of choice [21], this includes CaPs as well. On a cellular level, Sr ions have a dual mode of action: stimulation of osteoblasts and inhibition of osteoclasts[22]. Sr promotes formation of extracellular matrix (ECM) proteins produced by osteoblasts[23]. These effects might be useful in repair of the subchondral bone as well. In the available studies use of Sr containing biomaterials in bone defect repair is already well established [19, 24, 25] and it leads to improved or to at least unchanged new bone formation compared to the Sr-free groups [26]. However, the specific effects of Sr and even ACP on subchondral bone regeneration are still yet to be provided.

In particular, the combination of recently developed ACP with high specific surface area (>100 m^2^/g) [27–30] and Sr ions would provide excellent cues for ECM formation and subchondral bone tissue regeneration through sustaining of an ion-rich microenvironment. Upon contact with biological environment, dissolution of strontium, calcium and phosphate ion species is expected, which are favouring cues for ECM production and bone formation.

In this study, we modified the synthesis technology of ACP for incorporation of Sr, and developed a method to incorporate ACP/Sr-ACP granules into the Col/Col-Mg-HAp scaffold. Then we characterized physico-chemcical properties and the *in vitro* cytocompatibility of ACP or Sr-ACP granules and ACP/Sr-ACP enriched Col/Col-Mg-HAp scaffolds. To evaluate the osteogenic potential in osteochondral defects, we first investigated the biocompatibility and osteogenic effect of ACP/Sr-ACP enriched scaffold in an *in vivo* semi-orthotopic mouse model at the early phases of repair. Finally, the osteogenic effect of the Sr-ACP enriched Col/Col-Mg-HAp scaffold was investigated *in vivo* in a translational large animal (goat) osteochondral defect model.

## 2. Materials and methods

### 2.1 Synthesis of ACP and Sr-ACP

ACP and Sr-ACP granules used in the study were prepared from materials synthesized according to a wet precipitation technology developed previously [28]. Here, the synthesis technology was modified (use of calcium oxide instead of hydroxyapatite), and a novel synthesis procedure of ACP/Sr-ACP was developed as described further.

First, 2.71 g of CaO (calcined Ca(OH)_2_ (Jost Chemical Co., USA)) and 0.438 g of Sr(NO_3_)_2_ (Sigma-Aldrich, Germany) are mixed in deionized water (300 mL). The amount of Sr within Sr-ACP was chosen to be 50x the maximum amount reported of Sr in bone mineral (0.05 wt.% [31] i.e., 2.5 wt.%). The mixing was done with an overhead mixer MM-1000 (Biosan, Latvia) equipped with a propeller stirrer at 300-400 rpm at 20±2 °C. Then 14.48 mL of 2 M H_3_PO_4_ (75 %, “Latvijas Kimija” Ltd.) was admixed and the suspension was stirred for 30 min. Next 32.3 mL 3M HCl (Merck EMSURE^®^, Austria) at a rate of 5 mL/min was added. Resulting in dissolution of reagents, and thus a transparent solution containing calcium, phosphate, strontium, and nitrate ions was obtained. Next, after 30 min the mixing speed was increased to 450-550 rpm and an equimolar amount of 2M NaOH (Merck EMSURE^®^, Germany) was rapidly admixed to raise the pH and to induce precipitation of Sr-ACP. Then the stirring was continued for another 5 min until the reading of the pH electrode stabilizes (pH 10-11). Next, the precipitated Sr-ACP was separated by vacuum filtration. During the filtration, the Sr-ACP was washed with deionized water (1.5-2.0 L) to remove any formed water-soluble by-products, e.g., NaCl, from the precipitates. The presence of NaCl was tested by adding a few drops of 0.1 M silver nitrate to the solution that has passed the filter. When the formation of an opaque precipitate was not observed after the addition of the silver nitrate, it was considered that the solution does not contain NaCl. Then, the washed Sr-ACP was transferred onto glass Petri dishes, spread evenly, and dried at 80 °C for one hour in a drying oven with forced air circulation (UFE 400, Memmert, Germany). A schematic overview of the synthesis is shown in Figure 1. Synthesis of ACP was analogous but without the addition of Sr(NO_3_)_2_.

**Figure 1:**
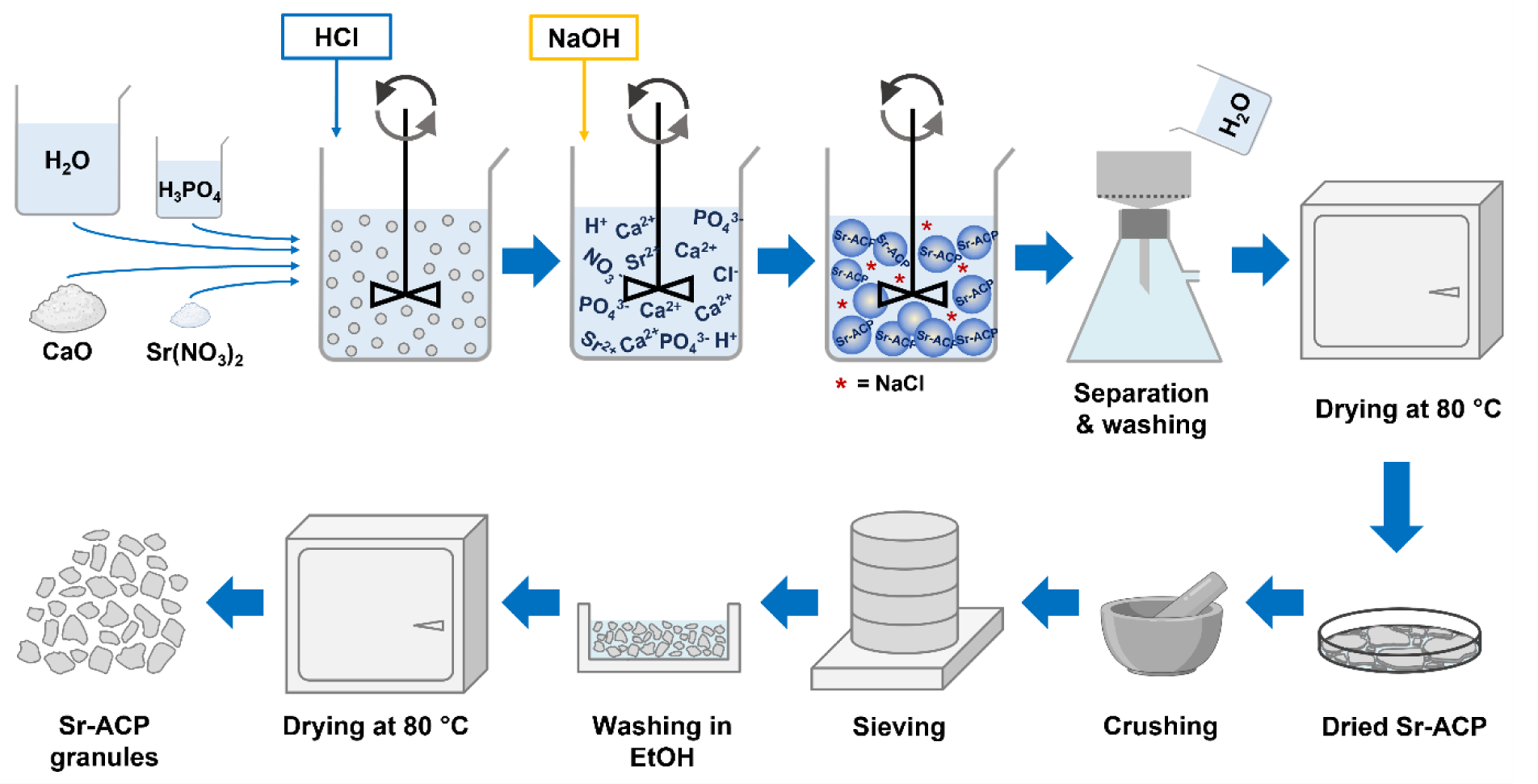
Schematic overview of Sr-ACP synthesis procedure (top) and dry granulation technology (bottom) for obtaining of Sr-ACP granules.

### 2.2 Production and characterization of ACP and Sr-ACP granules

ACP/Sr-ACP granules were manufactured using dry granulation technology (Figure 1) which involved milling of the synthesized ACP/Sr-ACP followed by sieving of the milled material to the desired range of granule size. In brief, the dried ACP/Sr-ACP precipitates were in the form of flat agglomerates (<3 mm thick). The agglomerates were manually crushed in a mortar and further sieved using sieves and a vibratory sieve shaker Analysette 3 (Fritsch GmbH, Germany). The sieving resulted in ACP/Sr-ACP granules in a size range of 100-150 µm. The debris that was formed during granulation was removed by rinsing the granules with ethanol (96 %). The rinsed granules were dried in a drying oven (UFE 400, Memmert, Germany) with forced air circulation at 80 °C (3 h). The manufactured ACP/Sr-ACP granules were characterized for their physicochemical properties as described below.

Phase composition of the synthesized ACP/Sr-ACP products was analysed using x-ray diffraction (XRD) with an X’Pert Pro (PANalaytical B.V., The Netherlands) diffractometer. The diffractometer was equipped with a Cu tube run at 40 kV and 30 mA. In the path of diffracted x-rays a Ni filter was installed to minimize Cu Kβ radiation. The XRD patterns were acquired in 2Theta range 10–70 ° with a step size of 0.0334 ° and time per step of 30.48 s. Powdered samples were put on a front-loading sample holder with a low background Si insert.

Information about chemical groups was gathered using a Fourier-transform infrared spectrometer (Varian 800 FT-IR, Scimitar Series, USA) in an attenuated reflectance (ATR, GladiATR^TM^, Pike technologies, USA) mode. Samples were finely ground and analysed in the form of a powder. FT-IR spectra were obtained at 4 cm^-1^ resolution co-adding 50 scans over a range of wavenumbers from 400 cm^-1^ to 4000 cm^-1^. Before each FT-IR measurement, a background spectrum was taken and later deducted from the sample spectrum.

Specific surface area (SSA) of the granules was determined by using an N_2_ adsorption system Quadrasorb SI Kr (Quantachrome Instruments, USA) with Autosorb Degasser AD-9 (Quantachrome Instruments, USA). Samples (0.5 g, n=3) were degassed at room temperature to remove any adsorbed volatiles. Calculation of the SSA was done according to Brunauer-Emmett-Teller (BET) theory. Next, the calculated particle size (d_BET_) was found using the following equation: d_BET_ = 6000/(SSA x density), assuming particles to be spherical.

Granule morphology was visualized using a field emission scanning electron microscope (SEM) Mira (Tescan, Czech Republic). SEM imaging was done at an accelerating voltage of 5 kV with both scanning electron (SE) and backscattered electron (BSE) detectors. Before the SEM imaging, samples were attached to sample holders with double-sided carbon tape and then coated with a layer of gold using sputter coater K550X (Quorum technologies, UK). Sputtering parameters were 25 mA for 180 s in an argon atmosphere with a sample rotation to obtain a homogenous coating.

Strontium concentration in Sr-ACP granules was determined using an inductively coupled plasma-optical emission spectrometry (ICP-OES, Thermo Scientific iCAP 7400, Waltham, MA, USA). The sample is dissolved in nitric acid (65 v/v%). The content (ppm) in the samples is determined by comparison with a predetermined standard curve. Sr (wt.%) was calculated on the basis of the sample weight.

### 2.3 Preparation and characterization of ACP/Sr-ACP granule containing collagen/collagen-magnesium-hydroxyapatite osteochondral scaffolds

Col/Col-Mg-HAp with/without ACP or Sr-ACP granules are biomimetic scaffolds that have a porous, 3-dimensional composite structure. The scaffold is composed of two layers: the cartilaginous layer consisting of Type I collagen and the bone layer consisting of a combination of Type I collagen (60%) and magnesium-hydroxyapatite (40%, Mg-HAp). Each layer of the scaffold is synthesised separately by a standardised process from an atelocollagen aqueous solution (1 wt.%) in acetic acid, isolated from equine tendon. The upper non-mineralised chondral layer of the scaffold is obtained by dissolving an acetic solution of Type I collagen in bi-distilled water by adding NaOH. The bone layer of the scaffold is obtained by nucleating nanostructured Mg-HAp into self-assembling collagen fibres, as occurs in the natural biological neo-ossification process. To stabilise the scaffold, the fibrous structures were chemically cross-linked for 16 hours at room temperature. After chemical cross-linking, ACP or Sr-ACP granules were added through a deposition by vacuum directly into the bone layer during the pre-filtration phase. The two layers are superimposed and afterwards they are freeze-dried. Finally, the scaffolds were gamma sterilized at 25 KGy.

ACP/Sr-ACP granule integration within the Col/Col-Mg-HAp scaffolds was evaluated using SEM/EDS and micro-CT techniques. Prior SEM imaging samples were cross sectioned with a scalpel. Further, the sample preparation procedure was the same as described above for ACP/Sr-ACP granules alone (section 2.2). Additionally, the scaffolds were analysed with an energy dispersive x-ray spectrometer (EDS) X-Max^N^ 150 (Oxford Instruments, UK) to obtain element distribution maps. To obtain element maps the electron gun was operated at 15 kV. Mapping area was selected by drawing a rectangle over the image of the sample. The total number of x-ray counts exceeded 10 000, thus ensuring good quality data. The EDS mapping was done with Inca software (Oxford Instruments, UK).

Further micro-CT analysis of the scaffolds was performed with a micro-CT 50 instrument (Scanco Medical, Switzerland). A sample holder with a diameter of 14 mm was used in which the scaffold was fixed with PU foam. Parameters of micro-CT control file were: energy 70 KV; intensity 114 µA; resolution - native; FOV 15.2; voxel size 3.4 µm; integration time 2004 s. Scans were done under a 0.5 mm thick Al filter. During image processing the sample was cut along the y and z axes. The samples were cut along the x axis when the PU foam had to be cut off.

### 2.4 *In vitro* cytotoxicity

To assess the possible cytotoxicity of the developed ACP/Sr-ACP granules and scaffolds, the *in vitro* cell viability was assessed. Granules or scaffolds were incubated in Dulbecco’s Modified Eagle Medium high glucose (DMEM, high glucose, Gibco, Waltham, MA, USA) supplemented with 10% fetal bovine serum (FBS, Gibco, Waltham, MA, USA) under gentle agitation for 24 hours at 37°C to obtain extracts. An extraction ratio of 0.2 g/ml for granules and 3 cm^2^/ml for scaffolds was considered, according to ISO 10993-12. Balb/c 3T3 clone A31 were seeded at 15000 cells /cm^2^ then incubated for 24 hours at 37°C before exposition to the extracts. Cells were incubated in culture medium with ACP or Sr-ACP extracts (25% and successive dilutions 15%, 8% and 2.5%) or scaffold extracts (100% and successive dilutions 40%, 16% and 6.4%) for 48 hours at 37°C in a humidified atmosphere with 5% CO_2_. Negative control (complete culture medium) and positive control for cytotoxicity (Phenol) were run in parallel. At the end of the incubation period, culture medium was removed and discarded. Cells were detached using trypsin solution. Then, a Trypan Blue solution with 10% FBS was added. Living cells were counted using a haemocytometer.

### 2.5 *In vivo* osteochondral defect mice model

To evaluate the biocompatibility and osteogenic capacity of ACP/Sr-ACP granules incorporated into the Col/Col-Mg-HAp scaffold *in vivo*, an osteochondral mouse model established by our group was used (Figure S1A) [32]. In order to model several larger critical sized bone defects in a mouse, we created a semi-orthotopic bovine bone defect model by implanting bovine bone plugs subcutaneously. Briefly, osteochondral defects (4 mm in diameter, 4 mm in depth) were created by a hand drill in bovine osteochondral biopsies (8 mm in diameter, 5 mm in height) harvested from metacarpal-phalangeal joints of 6 to 8 months old calves (LifeTec, Eindhoven, The Netherlands). The osteochondral explants were cultured overnight in alpha-Minimum Essential Medium (α-MEM; Gibco, Massachusetts, USA) supplemented with 10% fetal bovine serum (FBS, Gibco, Massachusetts, USA), 50 μg/mL gentamycin (Gibco, Massachusetts, USA), and 1.5 μg/mL fungizone (Gibco, Massachusetts, USA). Then the osteochondral defects were fitted with: (1) Col/Col-Mg-HAp scaffold-only (n=6, osteochondral scaffold, Finceramica, Italy, diameter: 4 mm, height: 4 mm), or (2) ACP enriched Col/Col-Mg-HAp scaffold (n=7), or (3) Sr-ACP enriched Col/Col-Mg-HAp scaffold (n=7). All osteochondral explants were covered with a circular 8 mm Neuro-Patch membrane (Braun, Melsungen, Germany) to prevent the ingrowth of host cells from the top.

Five 12-week-old NMRI-Fox1nu mice (Taconic, New York, USA) were randomly assigned and housed under specific-pathogen-free conditions with a regular day/night light cycle. Food and water were available ad libitum. The mice were allowed to adapt to the conditions of the animal facility for 7 days. The osteochondral explants were implanted subcutaneously on the back of the mice under 2.5-3% isoflurane anesthesia (1000 mg/g, Laboratorios Karizoo, Maharashtra, India). 4 osteochondral plugs were implanted in 4 pockets per mouse respectively. Staples (Fine Science Tools, Vancouver, Canada) were used to close the incisions and were removed 1 week after the implantation. To ensure pre- and post-operative analgesia, the mice received a subcutaneous injection of 0.05 mg/kg bodyweight of buprenorphine (Chr. Olesen & Co, Copenhagen, Denmark) 1 hour before surgery and 6-8 hour after surgery. Mice received a subcutaneous prophylactic antibiotic injection of 25 mg/kg body weight of Amoxicillin (Dopharma, Raamsdonksveer, Netherlands).

After 8 weeks, mice were euthanized by cervical dislocation under 2.5-3% isoflurane anesthesia and the osteochondral explants were harvested. All the samples were fixed in 4% formalin for 1 week for further processing. This animal experiment was approved by the Ethics Committee for Laboratory Animal Use (AVD101002016991; protocol #EMC 16-691-05).

### 2.6 *In vivo* translational large animal osteochondral defect model

A validated preclinical large animal bilateral osteochondral defect model was used to assess the osteogenic effect of the developed Sr-ACP enriched Col/Col-Mg-HAp scaffold. A gender balanced experimental unit of 12 skeletally mature Saanen goats (age: 2-3 years, weight: 35.8 ± 6.6 kg) was subjected to a bilateral arthrotomy under general anaesthesia as described before [33–35]. In short: all animals received a prophylactic antibiotic injection with amoxycillin clavulanic acid 8.75 mg/kg intramuscular (Noroclav, Norbrook, Ireland) and were intravenously sedated with butorphanol (0.2 mg/kg, Butador, Chanelle Pharma, Ireland) and diazepam (0.2 mg/kg, Diazemuls; Accord Healthcare, UK). A lumbosacral epidural block with lidocaine (2 mg/kg, Lidocaine HCl 2%, B. Braun Medical Inc., EU, Melsungen, Germany) and morphine (0.2 mg/kg, Morphine Sulphate 10 mg/mL, Kalceks, Latvia) was performed with the animal in sternal recumbency. Anaesthesia was induced with propofol IV to effect (max. 6 mg/kg, Propofol-Lipuro 1%, B. Braun Medical Inc., Melsungen, German) and was maintained with isoflurane (Vetflurane, Virbac Animal Health, Suffolk, UK) in 100% oxygen via a circle rebreathing system. All animals received analgesia with meloxicam IV (0.5 mg/kg, Rheumocam, Chanelle, Galway, Ireland); and morphine IV (0.2 mg/kg, Morphine sulphate, Mercury Pharmaceuticals, Dublin, Ireland) 90 min after the epidural block.

An arthrotomy of each stifle joint was performed in dorsal recumbency using a lateral parapatellar approach. Under constant irrigation with saline, a pointed 6 mm drill bit was used to drill an approximate 3-4 mm deep non-weight-bearing defect in the transition of the distal 1/3 to the middle 1/3 of the trochlear groove and in the central weight-bearing part of the medial femoral condyle. Subsequently, a custom-made flattened drill bit and a depth guide were used to create an exact flat 6 mm deep by 6 mm wide circular critical-sized osteochondral defect in a non-weight-bearing and a weight-bearing location. The joint was flushed with saline to remove any debris, and the defects were press fit with a similar-sized selected scaffold before surgical closure as described before. The left and right stifle joints of each goat were randomly assigned to one of the two treatment groups (within animal controlled) (Figure S1B): 1) Col/Col-Mg-HAp scaffold-only, and 2) Sr-ACP enriched Col/Col-Mg-HAp scaffold.

Following surgery, postoperative analgesia was provided (meloxicam 5 days) and goats were housed in indoor pens for daily postoperative welfare monitoring and scoring. Two weeks postoperatively, following the removal of skin sutures, animals were released to pasture or loose housing (weather dependent) for the remainder of the study period with daily health checks. An orthopaedic assessment (Table S1) was performed on the day of humane euthanasia under sedation with a barbiturate overdose at the predetermined endpoint at 6 months after surgery. Subsequently, all the joints, surrounding joint tissues, and synovial fluids were scored (Table S2), dissected, and photographed (Body Canon EOS R5, lens: Canon EF 100 mm f/2.8 L Macro IS USM, flash: Macro Ring lite MR-14EX II). Biopsies 1 cm by 1 cm square containing the entire osteochondral defects were harvested with an oscillating saw.

Ethical evaluation and approval were provided by the Health Products Regulatory Authority of Ireland (AE1898217/P142), the Animal Research Ethics Committee of University College Dublin (AREC-18–17-Brama) and the Lyons Animal Welfare Board (Health, Husbandry and Monitoring plans; 201907).

### 2.7 Macroscopic assessment of osteochondral defect repair

The quality of defect repair was assessed semi-quantitatively using the International Cartilage Repair Society (ICRS) macroscopic evaluation system (Table S3) [36] and a macroscopic scoring system (Table S4) developed by Goebel et al [37]. The ICRS scoring system rates cartilage repair tissue as Grade IV (severely abnormal), Grade III (abnormal), Grade II (nearly normal) or Grade I (normal). The Goebel Score describes articular cartilage repair with five major evaluation categories. The quality of defect repair was scored blinded on fresh samples by two independent assessors, and the scores were averaged for further calculation. All the samples were fixed in 4% formalin for 10 days after the macroscopic assessment.

### 2.8 Micro-computed tomography of subchondral bone defect repair

The harvested samples underwent micro-CT scans (Quantum GX, Perkin Elmer, USA) after fixation in 4% formalin *ex vivo*. For the bovine samples from the mouse model, the settings were: energy 90 KV, intensity 88 μA, 18 mm FOV, 36 μm isotropic voxel size. The micro-CT scan settings for goat samples were: energy 90 KV, intensity 88 μA, 36 mm FOV, 72 μm isotropic voxel size. All the scans were under an x-ray filter of Cu (thickness = 0.06 mm) and Al (thickness = 0.5 mm), and were calibrated using phantoms with a known density of 0.75 g/cm^3^, which were additionally scanned before and after each scan. A high-resolution mode was set, and a scan time of 4 minutes was used. Image processing included modest Gauss filtering (sigma=0.8 voxel, width=1 voxel) and segmentation using a single threshold. A cylindrical region (4 mm diameter and 5 mm height) in the defect was selected as a volume of interest (VOI). In this VOI the following morphometric parameters were measured: bone volume per total volume (BV/TV), trabecular thickness (Tb.Th), trabecular number (Tb.N), and trabecular separation (TB.Sp). Morphological analyses were performed using IPL (Scanco Medical AG, Bruettisellen, Switzerland).

### 2.9 Histology of osteochondral defect repair

After micro-CT scanning, the bovine osteochondral plugs from the mouse model were decalcified using 10% ethylenediaminetetraacetic acid (EDTA) for 4 weeks. The goat samples were decalcified for 3 weeks using 10% formic acid. Subsequently, all samples were embedded in paraffin and sectioned in 6 µm thin sections. Following dewaxing, H&E staining was performed with Hematoxylin (Sigma, Saint Louis, USA) and Eosin Y (Merck, Kenilworth, USA) to study general cell and tissue morphology. To visualize glycosaminoglycans in the extracellular matrix (ECM), dewaxed sections were stained with 0.1% Light green O (Fluka, Buchs, Switzerland) and 0.1% Safranin-O (Fluka, Buchs, Switzerland). To demonstrate the osteoclasts in the defects, Tartrate-resistant acid phosphatase (TRAP) staining was performed. Briefly, dewaxed sections were pre-incubated in sodium acetate (Sigma, Saint Louis, USA) and L(+) tartaric acid (Acros Organics, NJ, USA) buffer at room temperature for 20 minutes. Then naphtol AS-BI phosphate (Sigma, Saint Louis, USA) and fast red TR salt (Sigma, Saint Louis, USA) were added to the buffer and the slides were further incubated for 3 hours at 37 °C. To discriminate between calcified and non-calcified osteochondral tissue, RGB staining was performed using Alcian Blue (Sigma, Saint Louis, USA), Fast Green (Sigma, Saint Louis, USA), and Picrosirius Red (Sigma, Saint Louis, USA). NDP Software View2 (version 2.8.24, 2020 Hamamatsu Photonics K.K.) was used to measure the tissue volume in the osteochondral defect at three sections that were taken at the centre of the defect, and 0.5 mm and 1 mm from the centre for bovine samples from the mouse model or at the centre of the defect for the goat samples (Figure S2). The percentage of the defect covered with newly formed osteochondral tissue was calculated (Figure S3). Tissue volume in goat samples was independently measured by two investigators blinded to the experimental condition. The measurements of the two investigators were averaged for each section.

### 2.10 Statistical analysis

All statistical tests were performed using SPSS software 28.0 (SPSS inc., Chicago, USA). The repair tissue volume was expressed as mean ± standard deviation (SD). Multiple comparisons in cytotoxicity assessment, and between scaffold-only, ACP enriched scaffold and Sr-ACP enriched scaffold groups in bovine samples from the mouse model were analysed by a Kruskal-Wallis test. Statistically significant differences between the scaffold-only group and the Sr-ACP enriched scaffold group, or between trochlear groove and femoral condyle groups in goat samples were determined by Mann-Whitney U test. A p-value ≤ 0.05 was considered statistically significant.

## 3 Results

### 3.1 Characterization of ACP/Sr-ACP granules

The modified wet precipitation technology successfully yielded ACP and Sr-ACP materials. An overview of ACP/Sr-ACP granule physicochemical characteristics is given in Table 1 and Figure 2.

**Figure 2.**
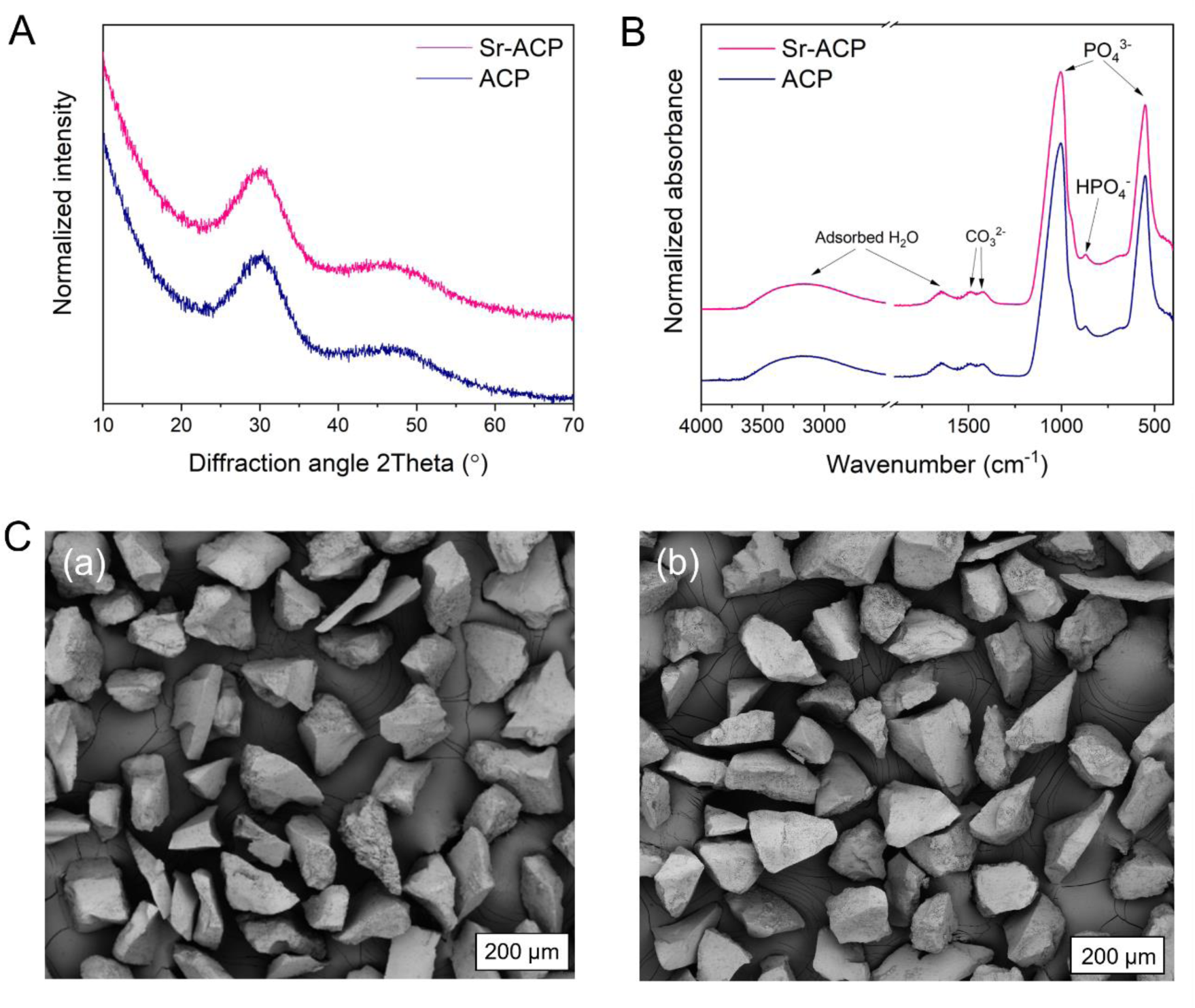
Physicochemical characteristics of ACP and Sr-ACP granules: (A) XRD patterns showing wide diffraction maxima characteristic to ACP materials, (B) FT-IR spectra demonstrating chemical group information and the hydrated nature of both materials, and (C) SEM images of ACP (a) and Sr-ACP (b) irregularly shaped granules.

**Table 1.** Values of Sr concentration, specific surface area (SSA), and calculated particle size d_BET_ for ACP and Sr-ACP granules.

| | Sr conc., wt.% | SSA, m<sup>2</sup>/g | $d_{\text{BET}}$, nm |
| --- | --- | --- | --- |
| ACP | 0.01 | $113 \pm 2$ | 21 |
| Sr-ACP | $2.49 \pm 0.04$ | $115 \pm 2$ | 20 |

The XRD patterns confirmed the amorphous character of the obtained products (Figure 2A). The experimental Sr concentration of Sr-ACP (Table 1) was determined with ICP-MS at 2.49 ± 0.04 wt.% (n=3), which compared well with the theoretical value of 2.5 wt.%. The FT-IR spectra (Figure 2B) demonstrated the hydrated and carbonated nature both of ACP and Sr-ACP. Introduction of Sr ions in the given concentration did not reveal any structural changes that could be observed with XRD and FT-IR. The specific surface area of both ACP and Sr-ACP granules was high (> 100 m^2^/g) with particle size d_BET_ being 20-21 nm (Table 1). The dry granulation technology produced irregular shape granules with sharp edges (Figure 2C). The sharp edges of the granules originate from the milling of the ACP agglomerates. Granule surfaces at the macro level are smooth and non-porous. By measuring granule dimensions from the SEM images, an average value of the experimental granule size was determined to be 187 ± 35 μm.

The final step of the granule production was granule washing with ethanol to remove any debris that may have originated from the granulation process. To assess whether the rinsing procedure has an impact on the structure of the ACP materials, granules were characterized with FT-IR (Figure S4). No differences in FT-IR spectra of ACP granules before and after the rinsing with ethanol were detected.

Before *in vitro* and *in vivo* experiments, materials must be sterilized; in this study, gamma irradiation was used. To ensure amorphous granule composition remained unaffected post-sterilization, phase and chemical composition were analyzed using XRD and FT-IR. Obtained results demonstrated that gamma irradiation sterilization of ACP granules was effective, with no detectable changes in composition or crystallinity (Figure S5).

### 3.2 ACP/Sr-ACP granule containing Col/Col-Mg-HAp scaffolds

Addition of ACP/Sr-ACP granules to the Col/Col-Mg-HAp scaffold is an additional step for the manufacturing process of the scaffolds. The newly developed ACP/Sr-ACP granule containing Col/Col-Mg-HAp scaffolds were examined with three-dimensional micro-CT analyses to assess granule 3D distribution within the ACP granule containing scaffolds. The ACP granules are well and homogeneously distributed in the bottom layer of scaffold (Figure 3A). The SEM image (Figure 3B) shows the bilayered structure of the scaffold: collagen-only layer on top and collagen/Mg-HAp/ACP granule layer on the bottom. Both layers of the freeze-dried scaffold have a porous structure, which is governed by collagen. In the bottom layer the incorporated micron-sized ACP granules can be seen, while the nanoparticles of Mg-HAp can’t be visualized at given magnification. SEM-EDS element maps of Ca, P, C, and Mg of collagen/Mg-HAp/ACP granule scaffolds are shown (Figure 3C). Element maps of the same area (Figure 3C) confirm presence of the ACP granules as well. As Ca and P are the main constituents of ACP, the high contrast areas in Ca and P element maps matches ACP granule placement in the SEM image (Figure 3B). EDS map of Mg designates the location of the biomimetically deposited Mg-HAp nanoparticles onto the fibers of the collagen scaffold’s bottom layer. Map of C demonstrates the presence of collagen throughout the mapped area; higher intensity area of C is visible for the top layer which contains only collagen and no calcium phosphates. SEM inspection of ACP granule containing scaffolds showed that the granules have a good compatibility with the scaffold’s main component - collagen. SEM images (Figure 3D), showed that the ACP granules were incorporated in the collagen fibers of the scaffold. Collagen fibers were attached to the surface of the granule and stretch across it.

**Figure 3.**
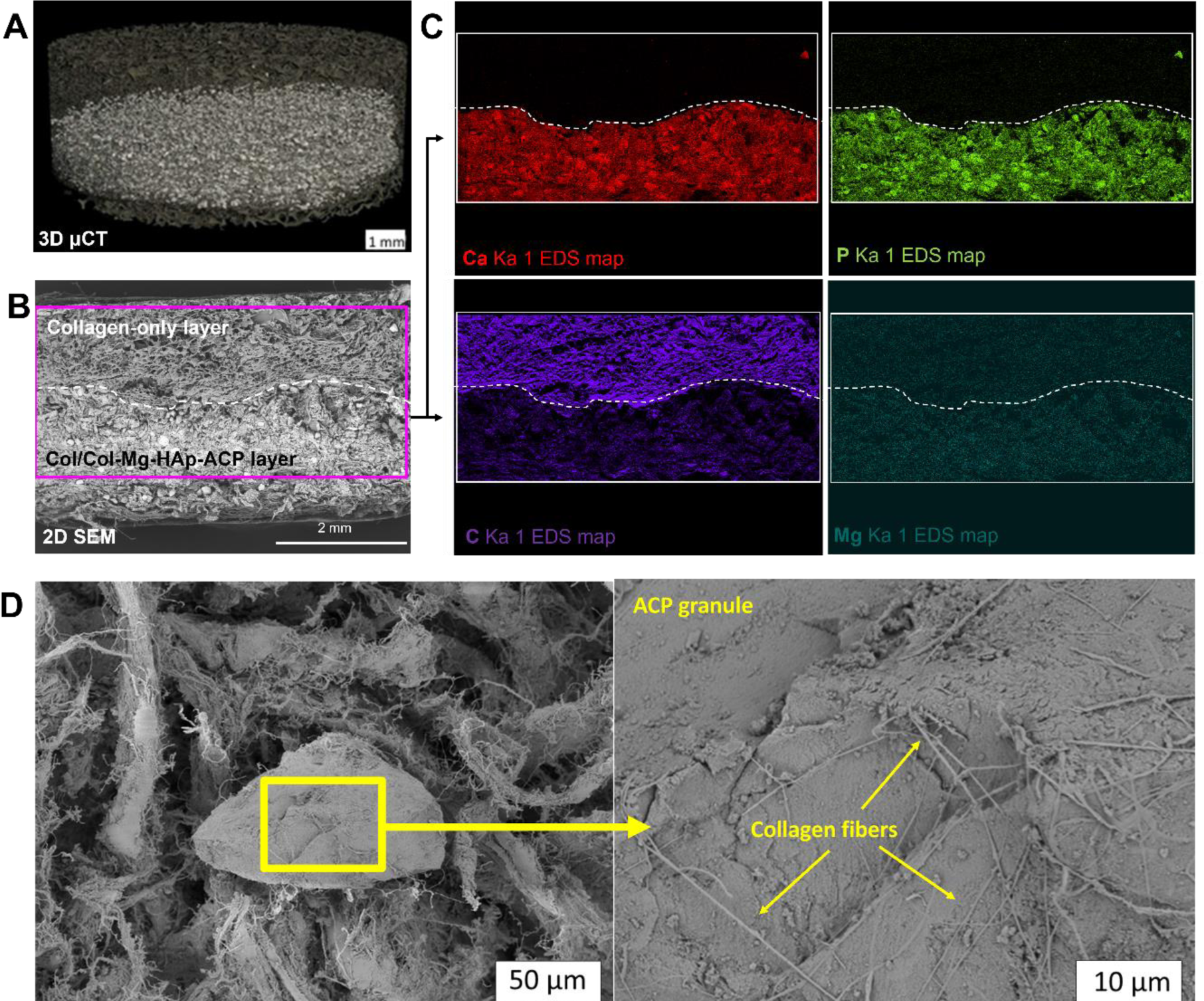
Representative micro-CT image (A) of Sr-ACP granule containing Col/Col-Mg-HAp scaffold. SEM image in backscattered electron (BSE) detector mode (B) of cross-section of the ACP granule containing Col/Col-Mg-HAp scaffold, where the top layer is collagen and the bottom layer is collagen-Mg-HAp layer enriched with ACP granules, and where the drawn rectangle shows EDS mapping area. EDS element (Ca - red, P - green, Mg - dark green, C - purple) maps (C) of the scaffold visualized on (B), where the dashed line shows the border between the both layers and the brightest areas in Ca and P maps represent positions of the ACP granules (D) SEM images of ACP granule containing scaffold where single ACP granule (left) and close-up view of the surface of the ACP granule covered in collagen fibers (right) is shown.

### 3.3 Cytotoxicity assessment

The 25%, 15%, 8% and 2.5% dilutions of extracts that was harvested after 24 hours incubation of ACP or Sr-ACP (2.49 wt.% Sr) granules were not cytotoxic (Figure 4A). 8% and 2.5% extract dilutions equivalent to 30% and 10% in weight of the scaffold, respectively. Therefore, enriched scaffold with 30 wt.% ACP or Sr-ACP granules were prepared for cytotoxicity assessment. Extracts from the ACP or Sr-ACP (30 wt.%) enriched scaffolds were cytotoxic for 100% and 40% extracts, and reached a non-cytotoxic level from dilution 16% (Figure 4B).

**Figure 4.**
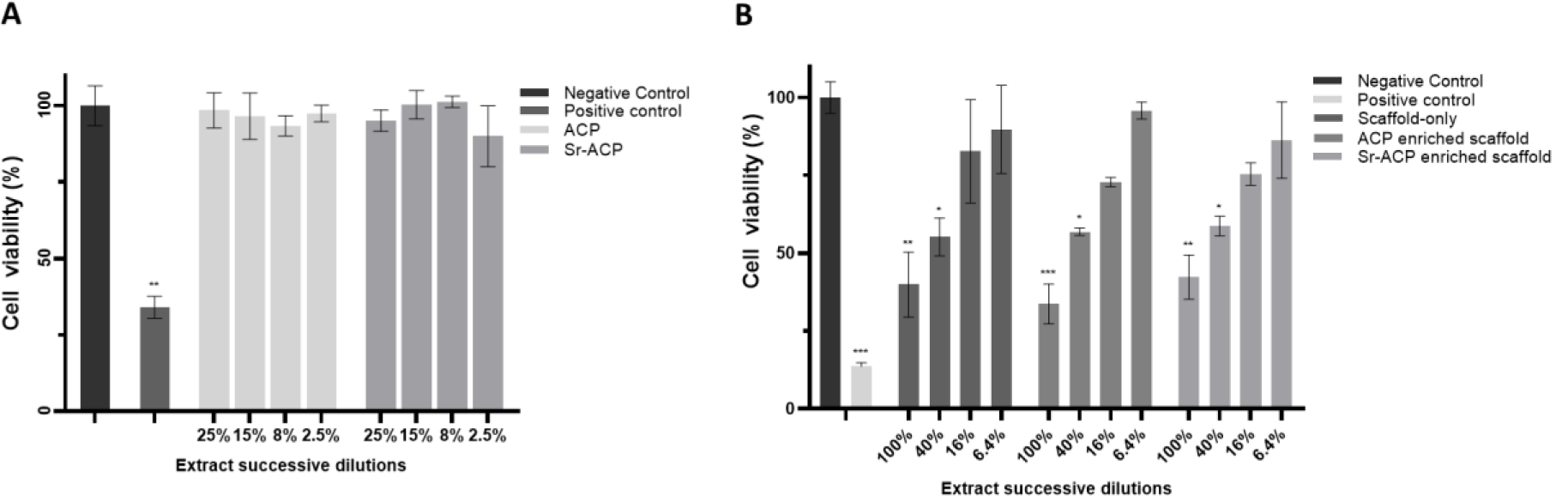
Cytocompatibility evaluation of Balb/c 3T3 clone A31 cells exposed to ACP/Sr-ACP granules extraction (A, n=4) and ACP/Sr-ACP enriched scaffold extraction (B, n=3). Cell viability (%) is the ratio of test condition and negative control. *** P<0.001, ** P<0.01, * P<0.05 compared to negative control.

### 3.4 Effect of ACP and Sr-ACP on osteochondral defect repair in an *in vivo* mouse subcutaneous model

An *in vivo* early osteochondral repair phase semi-orthotopic mouse model was used to assess the effect of ACP or Sr-ACP incorporation into the Col/Col-Mg-Hap scaffold. After 8 weeks, remnants of the collagen-only layer were observed in the cartilage region of defects, while the Col-Mg-HAp layer in the subchondral bone defect was mostly degraded and replaced by bone-like tissue (Figure 5A). Notably, ACP or Sr-ACP granules can still be seen after 8 weeks, and were well distributed in the newly formed tissues (Figure 5A). Some osteoclasts attaching to the granules in the bone tissue were demonstrated by TRAP staining (Figure 5A). The subchondral bone defects were filled with newly formed osteochondral tissue, indicating a good biocompatibility of ACP and Sr-ACP granules. Slightly more osteochondral repair tissue was found in the osteochondral defects loaded with Sr-ACP enriched scaffolds (89.3 ± 7.2%) compared to the scaffold-only (87.2 ± 11.1%) or ACP enriched scaffolds (80.2 ± 21.5%), albeit no significant differences in tissue volumes were found (Figure 5B).

**Figure 5.**
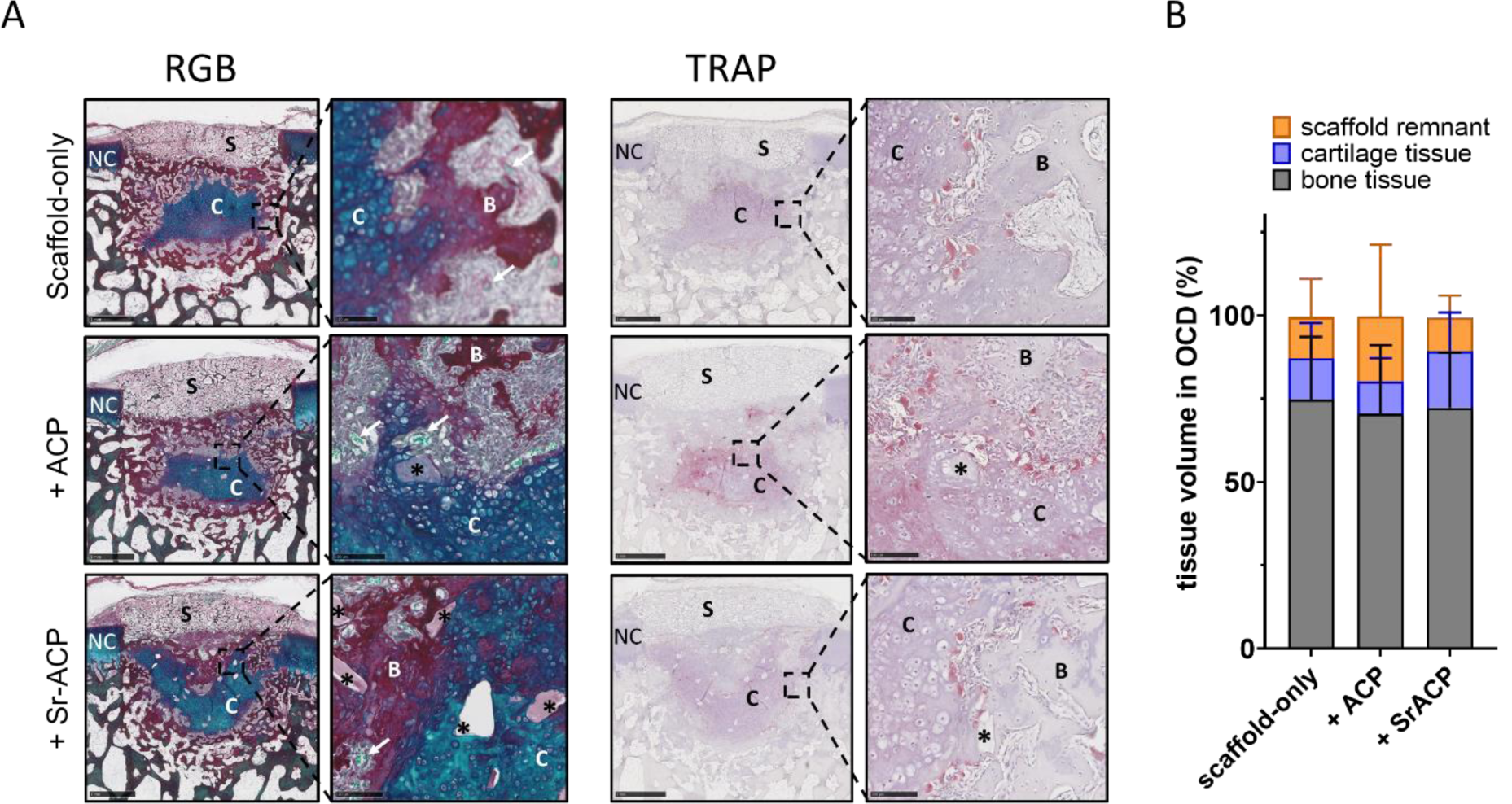
ACP and Sr-ACP showed a good biocompatibility for osteochondral repair *in vivo*. (A) Representative images of the 8-week repair constructs stained with RGB (Alcian Blue, Fast Green, and Picrosirius Red) and immunohistochemical staining for Tartrate-Resistant Acid Phosphatase (TRAP). Scale bars indicate 1 mm and 100 µm, respectively. NC: native cartilage; C: newly formed cartilage-like tissue; B: newly formed bone-like tissue; S: remnants of the scaffolds; *: ACP or Sr-ACP granules. (B) The percentage of tissue volume calculated in the osteochondral defects (OCD).

### 3.5 Effect of Sr-ACP on osteochondral defect repair in an *in vivo* large animal translational model

#### 3.5.1 Clinical observations and scaffold implantation

The Sr-ACP enriched Col/Col-Mg-HAp scaffold demonstrated good repair in the mouse model and therefore the osteogenic capacity of Sr-ACP granules was further investigated in a bilateral validated translational goat osteochondral defect model in the knee. Scaffolds were successfully implanted into osteochondral defects created in the trochlear groove (a non-weight-bearing location) and the medial femoral condyle (a weight-bearing location) of both knees. All animals recovered well postoperatively except for one goat that died 2 weeks post-surgery due to clostridium disease unrelated to the surgery or the experiment. The macroscopic appearance 2 weeks post-surgery showed that scaffolds were stable at both medial femoral condyle and trochlear groove osteochondral defect sites (Figure S6A). The two layers of the scaffold can clearly be seen in the osteochondral defects histologically at two weeks (Figure S6B). Another two goats died 4 months post-surgery, and one goat died 5 months post-surgery, again caused by clostridium disease despite vaccination and due to reasons unrelated to the surgery and the experiment. The remaining eight goats remained in good health throughout the study. At the predetermined 6-month endpoint the orthopedic exam demonstrated normal locomotion and excellent joint mobility in all goats.

All the joints, surrounding joint tissues, and synovial fluid were scored macroscopically on opening of the joints. There was no evidence of inflammatory responses or construct delamination in the treated joints at the time of retrieval. No joint swelling, effusion, mobility abnormalities or adhesions were found. Synovial fluid and membrane were normal and no indications of patellar instability/luxation were found.

#### 3.4.2 Tissue repair in the osteochondral defects

The samples from the goats that unexpectedly died after 4-5 months, revealed that the scaffolds had been degraded completely, and the osteochondral defects were mostly filled with repair tissue histologically (Figure S7). Overall, at 6 months, well-structured subchondral trabecular bone was observed in most trochlear groove and femoral condyle subchondral bone defects demonstrated by reconstructed micro-CT images, macroscopic sectional view and histology (Figure 6, 7). Reconstructed subchondral bone defect images showed an area with no trabecular bone either underneath or at the bottom of the defects unrelated to the defect location or the scaffold type (Figure 6A, 7A). Histological images demonstrated that these areas found in micro-CT images were filled with bone marrow and were dissimilar to cysts (Figure 6C, 7C).

**Figure 6.**
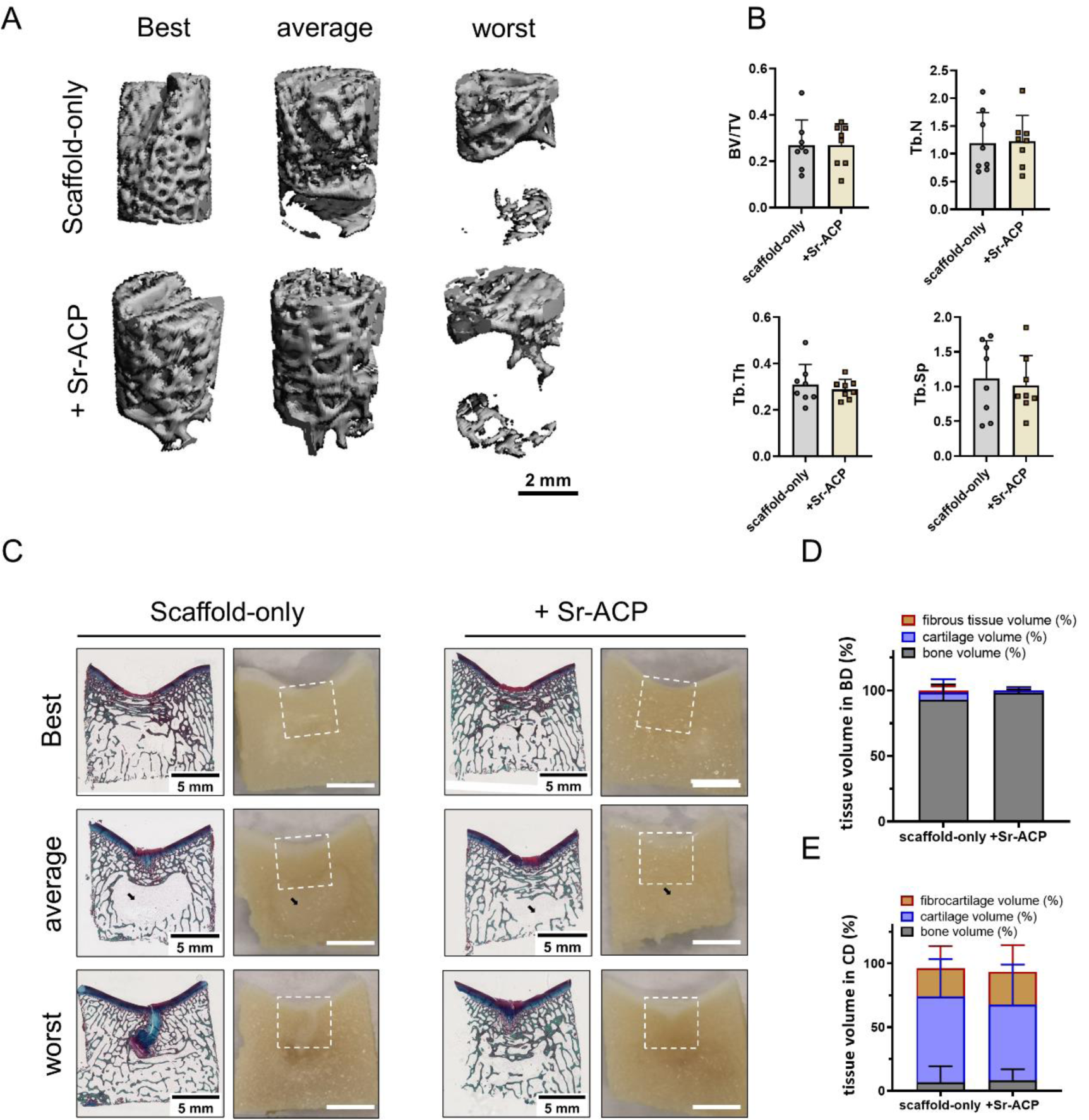
Tissue repair in the trochlear groove defect sites. (A) Representative micro-CT reconstructions treated with either scaffold-only or Sr-ACP enriched scaffold. Best, average, and worst repaired samples are presented. The scale bar indicated 2 mm. (B) BV/TV, trabecular thickness (Tb. Th [mm]), trabecular number (Tb. N [1/mm]), and trabecular separation (Tb. Sp [mm]) in the bone defects after 6 months. (C) RGB (Alcian Blue, Fast Green, and Picrosirius Red) staining and macroscopically sectional view of osteochondral defects treated with either scaffold-only or Sr-ACP enriched scaffold. Best, average, and worst repaired samples are presented. White squares indicated 6*6 mm osteochondral defects. Black arrows indicated the structure with only bone marrow. The scale bar indicated 5 mm. (D) The percentage of tissue volume calculated in the subchondral bone defects (BD). (E) The percentage of tissue volume calculated in the cartilage defects (CD).

**Figure 7.**
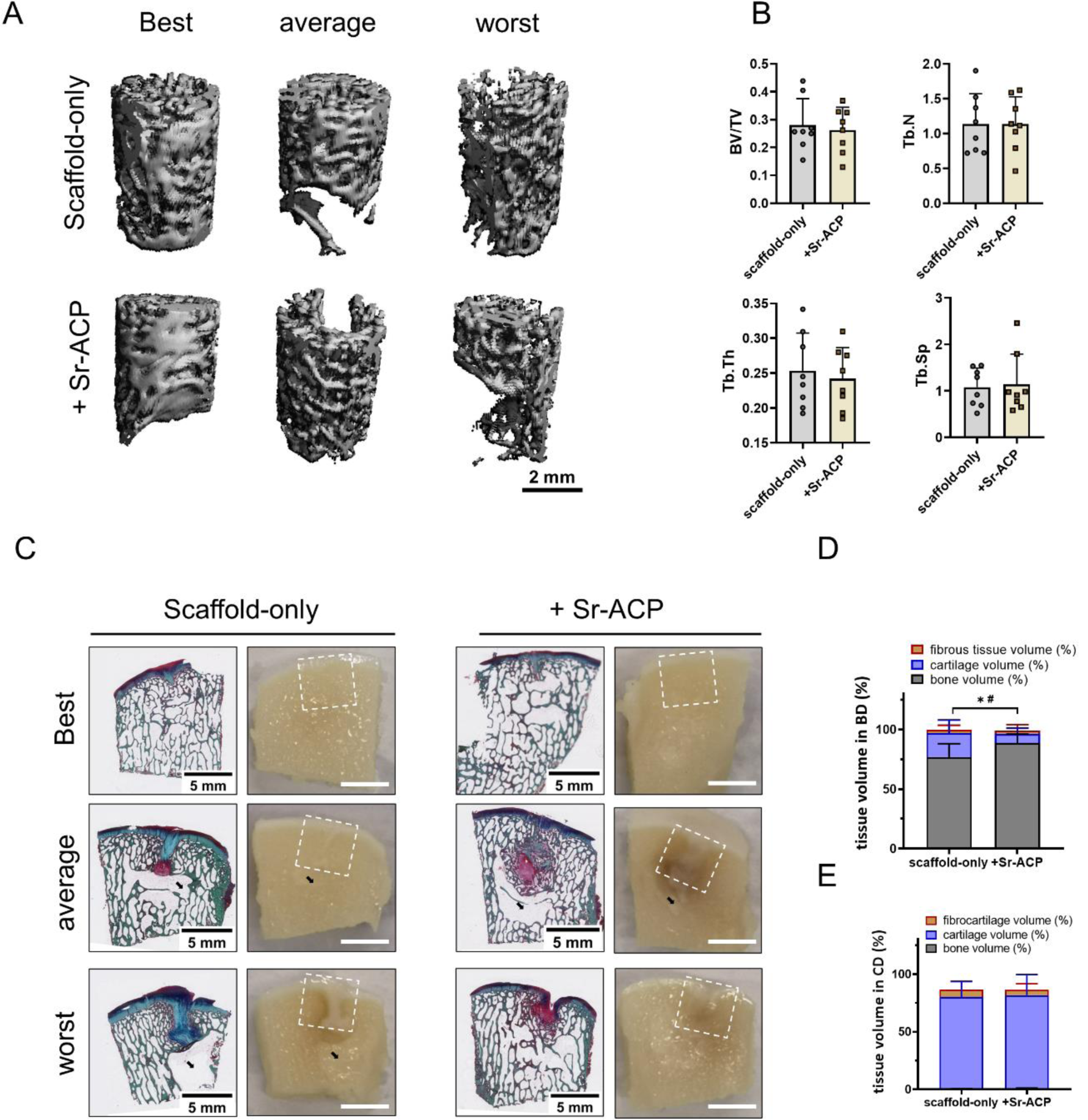
Tissue repair in the femoral condyle defect sites. (A) Representative micro-CT reconstructions treated with either scaffold-only or Sr-ACP enriched scaffold. Best, average, and worst repaired samples are presented. The scale bar indicated 2 mm. (B) BV/TV, trabecular thickness (Tb. Th [mm]), trabecular number (Tb. N [1/mm]), and trabecular separation (Tb. Sp [mm]) in the subchondral bone defects after 6 months. (C) RGB (Alcian Blue, Fast Green, and Picrosirius Red) staining and macroscopic images of osteochondral defects treated with either scaffold-only or Sr-ACP enriched scaffold. Best, average, and worst repaired samples are presented. White squares indicated 6*6 mm osteochondral defects. Black arrows indicated the structure with only bone marrow. The scale bar indicated 5 mm. (D) The percentage of tissue volume calculated in the subchondral bone defects (BD). * P<0.05 in cartilage-like tissue, # P<0.05 in bone-like tissue. (E) The percentage of tissue volume calculated in the cartilage defects (CD).

In the non-weight-bearing trochlear groove location reconstructed micro-CT images showed no significant difference in the BV/TV, Tb. Th, Tb. N and Tb. Sp in subchondral bone defects (Figure 6A, B). The macroscopic cross-sectional view and histology further confirmed the well-repaired subchondral bone (Figure 6C). Bone-like tissue (including the bone marrow) was quantified on RGB stained histology. After 6 months, slightly more bone tissue (98.0 ± 29.0% vs. 92.7 ± 11.9%, P=0.328) was found in the subchondral bone defects when the Sr-ACP was incorporated into the scaffolds compared to the scaffold-only, although no statistically significant difference was found (Figure 6D).

In the weight-bearing femoral condyle location no significant difference in the BV/TV, Tb. Th, Tb. N, and Tb. Sp was observed at 6 months (Fig 7A, B). Overall, 96.9 ± 3.8%

(scaffold-only group) and 96.0 ± 5.6% (Sr-ACP enriched scaffold) of the subchondral bone defects were filled with osteochondral tissue (Fig 7D). However, when looking specifically at the study target, the bone layer of the osteochondral unit, significantly more bone tissue was found (P=0.038, Fig 7D) in the subchondral defects loaded with Sr-ACP enriched scaffold (88.6 + 7.6%) compared to scaffold-only (76.7 ± 11.4%). Interestingly, significantly more bone-like tissue was regenerated in the trochlear groove subchondral bone defect sites compared to the medial femoral condyle subchondral bone defect sites when scaffold-only (92.7 ± 11.9% vs. 76.7 ± 11.4%, P=0.007) or Sr-ACP enriched scaffolds (bone-like tissue: 98.0 ± 2.9% vs. 88.6 ± 7.6%, P=0.007) were implanted in the osteochondral defects.

The cartilage part of the defects treated with either scaffold repaired well with good integration into the surrounding native tissue macroscopically at 6 months post-implantation (Figure S8A, B). Only small, scattered fissures or cracks were observed on some surfaces of the defects and no noticeable depressions were observed overall. In trochlear groove defects, the macroscopic ICRS and Goebel scores for the scaffold-only group had a median score of 10.19 ± 1.65 out of 12 and 17.19 ± 3.39 out of 20, respectively (Figure S8C). All the samples were classified as normal (grade I) or nearly normal (grade II) cartilage except for one sample (grade III). For the Sr-ACP enriched scaffold group, the macroscopic ICRS and Goebel scores were 9.50 ± 2.98 and 16.63 ± 3.93, respectively (Figure S8C). Two defects repaired with the Sr-ACP enriched scaffold were classified as abnormal (grade III). Macroscopic assessment of femoral condyle defects repaired with the scaffold-only resulted in median ICRS scores of 10.13 ± 0.83, and median Goebel scores of 18.69 ± 0.37 at 6 months (Figure S8D). The defects fitted with the Sr-ACP enriched scaffold were scored median ICRS scores of 9.94 ± 1.27, and median Goebel scores of 18.56 ± 1.02 (Figure S8D). All the samples were classified as nearly normal (grade II) cartilage. Overall, no significant difference was observed in cartilage repair between these two conditions with both scoring systems. Histologically, cells with a rounded morphology within the cartilage region were found residing within lacunae and with alignment typical of native cartilage. Both scaffolds demonstrated cartilaginous tissue formation by positive GAG and collagen RGB staining (Figure 6C, 7C). No significant difference could be found between the scaffolds (Figure 6E, 7E).

## 4. Discussion

The main finding of this study is that the addition of Sr-ACP granules into a clinically used osteochondral scaffold is a feasible and effective strategy to improve its bone repair capacity in *in vivo* osteochondral defects. The subcutaneous mouse osteochondral defect model demonstrated good biocompatibility and an overall good early tissue response of both ACP and Sr-ACP enriched Col/Col-Mg-HAp scaffolds, whereas a better bone formation was obtained in subchondral bone defects treated with the Sr-ACP enriched scaffolds in particular in the weight-bearing femoral condyle subchondral bone defect at 6 months in a goat model.

The new strategy proposed in this study is based on the modification of a clinically used Col/Col-Mg-HAp scaffold through the incorporation of ACP or Sr-ACP granules with a high specific surface area (>100 m^2^/g) and a hydrated and carbonated nature. A simple, fast, cost-effective, and scalable method for the preparation of ACP was used in this study. ACP or Sr-ACP granules with a large specific surface area and hydrated and carbonated nature were well distributed in the Col/Col-Mg-HAp scaffold. Due to the potent effects of calcium and phosphate ions on bone cells, and their presence in large quantities in bone tissue, calcium phosphates (CaPs) are of high interest in the bone repair biomaterial field [38]. In nature, ACP is involved in the early stages of bone mineralization [39] and the formation of complex CaP structures during bone mineral shaping and structuring [38, 40]. Previous studies on ACP have demonstrated excellent biocompatibility and bioactivity of this product *in vitro* [41] as well as good biodegradability, osteoconductivity, and osteogenic potential also in *in vivo* osteochondral defect models [42, 43]. On the other hand, the main inorganic component of bone is low crystalline apatite that highly resembles the chemical structure of HAp [44–46]. The addition of HAp into the bone layer can further improve the osteogenic potential of a collagen-based scaffold *in vivo* [47–51]. Therefore, the biphasic combination of ACP and HAp materials was expected to improve bone regeneration in an osteochondral defect. High crystallinity and stoichiometry of HAp lead to rather slow rates of dissolution, thereby improving mechanical properties of the scaffold and long-term bone regeneration [52]. ACP, in the meantime, can act in the early stages of biomaterial remodelling favouring the onset of bone deposition due to its high solubility and amorphous structure [40].

In the meantime, we have successfully combined an alternative local Sr^2+^ delivery carrier in the form of ACP granules within the Col/Col-Mg-HAp scaffold to further improve the bone regeneration. Sr and Ca are chemically very similar in ion size and have the same charge (+2) [53], thus Sr incorporation in calcium rich materials can be achieved. The majority of *in vitro* studies support a dual effect of Sr^2+^ on bone tissue: 1) stimulating bone formation by increasing proliferation and differentiation of osteoblasts, and inhibiting their apoptosis [21, 54–56]; 2) hindering bone resorption by inhibiting the formation and differentiation of osteoclasts and promoting their apoptosis [55–57]. Our *in vivo* mouse study showed slightly more osteochondral repair tissue in the defects loaded with Sr-ACP enriched scaffolds compared to ACP enriched scaffolds, although in both conditions, osteoclasts attaching to the ACP or Sr-ACP granules were observed.

The possible structural transformation of ACP into other calcium phosphate compounds raised problems for mass production, processing and storage [40]. The synthesis route for the preparation of amorphous calcium phosphates we used in this study enabled stability of ACP in air in a dried state for at least 7 months [28]. Trace amounts of various ions have been corroborated to affect ACP transformation [58, 59]. Mg^2+^ is an effective inhibitor for the ACP phase transformation by changing the internal structure of ACP and reducing solubility [60–62]. Furthermore, Sr^2+^ can stabilize ACP as well [58]. Interestingly, the presence of Sr^2+^ was reported to significantly enhance the stabilization effect of Mg^2+^ on ACP due to a synergic effect, which might be due to that Sr^2+^ promotes the exclusion of Mg during HAp nucleation from ACP [59]. Therefore, a relatively stable phase of ACP was expected in a Col/Col-Mg-HAp-Sr-ACP scaffold before implantation. After implantation Sr-ACP/ACP granules eventually would transform into poorly crystalline calcium phosphate phase resembling bone mineral. Our *in vivo* study demonstrated that incorporated granules were still present after 8 weeks in mice, and were degraded after 4 months in goats, when there was already sufficient bone regeneration, although the composition (Sr-ACP/ACP granules or calcium phosphate phase) of the granules found on the histology was not confirmed.

Here, the Col/Col-Mg-Hap scaffolds modified with ACP and Sr-ACP were investigated on the sequential use for osteochondral defects in *in vivo* models, from a small animal model to a large animal model. These two models, used together, allowed us to investigate the possible effect of incorporating ACP or Sr-ACP into the Col/Col-Mg-HAp scaffold used for osteochondral repair and to bring our approach a step closer to the physiological and mechanical conditions in the human osteochondral environment. We first confirmed biocompatibility and osteogenic properties of the modified Col/Col-Mg-HAp scaffolds in the mouse model as the first screening. The semi-orthotopic model allows a minimally invasive surgery and a multiple graft testing possibility [63], in line with the increasing ethical requirements on animal experiments. The results showed that, after 8 weeks, the bone-like layer in the subchondral bone defect was mostly degraded and replaced by bone-like tissue. The presence of repaired bone tissue together with the lack of side effects in all the experimental groups demonstrated a safe and good repair capacity of both ACP and Sr-ACP enriched scaffolds. In fact, both the native osteochondral Col/Col-Mg-HAp scaffold and the incorporated inorganic granules have been shown to be biocompatible and biodegradable [41-43, 49, 64, 65]. The three treatment groups showed the presence of repair tissue, with no significant differences among the different scaffolds. This indicates, on one side, that the granule insertion did not interfere with the healing process and, on the other side, that the overall healing process was very fast. This was expected, given the native repair capacity of the Col/Col-Mg-HAp scaffold per se, as supported by clinical evidence [9] together with the intrinsic nature of osteochondral defects in smaller animals, which tend to heal quickly if compared with larger animals [66]. Consequently, 8 weeks represent a relatively late time point in this model. Therefore, considering the quick repair response, and also the lack of synovial fluid, mechanical loading and complete immune system in the mouse [63], the use of a more advanced translational large animal model, suitable for comparison with human conditions, was a logical subsequent step. Thus, after the preliminary evaluation in mice, the most promising scaffold modification, the addition of Sr-ACP granules, was selected to be tested in a goat orthotopic osteochondral defect model.

The goat model is a fully immune competent model using outbreed animals, and offers advantages regarding joint size, cartilage and subchondral bone thickness, accessibility for arthroscopic procedures, and limited intrinsic healing capacity[67]. Also, it can provide the opportunity to assess tissue regeneration in two different mechanical loading environments within the same joint. In particular, the Col/Col-Mg-HAp and the Sr-ACP enriched Col/Col-Mg-HAp scaffolds were successfully implanted in the trochlear groove, with no/low direct mechanical loading, and in the medial femoral condyle, with direct mainly compressional mechanical loading [68]. In this goat model, significantly more bone was regenerated after 6 months in the subchondral bone defects of the biomechanically more challenging femoral condyle lesions when Sr-ACP was incorporated into the scaffold compared to scaffold-only. In fact, during its metabolism, bone incorporates and releases various trace elements (Na, Mg, Sr, Zn, Si etc.) into the cellular microenvironment [53]. Similar element/ion release in the cellular microenvironment was expected when Sr-ACP was incorporated into the Col/Col-Mg-HAp scaffold, where Sr^2+^, Mg^2+^, Ca^2+^, and PO_4_^3-^ should be released from the scaffold during the healing process. Although no significant difference in bone formation was found in mice treated with scaffolds modified with ACP with and without Sr. The overall good osteochondral regeneration obtained with the scaffold-only may have hindered the possibility to detect a significant improvement in this model, which did not show the same criticalities observed in terms of osteochondral regeneration in humans. In the more challenging and translational goat model, the incorporation of Sr-ACP into the scaffold was significantly more effective in regenerating bone tissue compared to the scaffold-only, as shown by the histological analysis. Overall, the scaffold-only and Sr-ACP enriched scaffolds regenerated a similar volume of osteochondral tissues, which means more cartilage-like tissue was present in the subchondral bone defect with the scaffold-only. These cartilage-like tissues might be ossified afterwards. In other words, there might be an acceleration effect of Sr-ACP at the earlier stage of repair. However, in this study, bone repair at only one time-point was assessed in the goat model. Therefore, the early cellular responsiveness that leads to a potential acceleration at this stage of repair, or long-term osteogenesis, which is known to end within 10 to 12 months [69], was not investigated. An effect could have been missed at its full extent by having a study focus of 6 months. This may also explain how, unlike what was observed by the histological analysis, no significant difference in bone volume was found by micro-CT analysis.

## 5. Conclusion

The incorporation of osteogenic Sr-ACP granules showed to be a feasible and promising strategy, as it improves the bone formation capacity of a Col/Col-Mg-HAp scaffold in subchondral bone defect repair. We propose that the use of Sr-ACP granules in the bone layer of a bilayered osteochondral scaffold would lead to enhanced osteochondral defect repair.

## Author statement

Jietao Xu: Data curation, formal analysis, investigation, writing-original draft, review and editing. Jana Vecstaudža: Data curation, formal analysis, investigation, writing-original draft, review and editing. Marinus A. Wesdorp: Investigation, writing-review and editing. Margot Labberté: Data curation, investigation, writing-review and editing. Nicole Kops: Investigation, writing-review and editing. Manuela Salerno: Writing-original draft, review and editing. Joeri Kok: Data curation, formal analysis, writing-review and editing. Marina Simon: Data curation, formal analysis, writing-review and editing. Marie-Françoise Harmand: Data curation, formal analysis, writing-review and editing. Karin Vancíková: Investigation, writing-review and editing. Bert van Rietbergen: Writing-review and editing. Massimiliano Maraglino Misciagna: Resources, writing-review and editing. Laura Dolcini: Resources, writing-review and editing. Giuseppe Filardo: Conceptualization, funding acquisition, project administration, writing-review and editing. Eric Farrell: Conceptualization, funding acquisition, supervision, writing-review and editing. Gerjo J.V.M. van Osch: Conceptualization, data curation, funding acquisition, project administration, supervision, writing-original draft, review and editing. Jānis Ločs: Conceptualization, data curation, funding acquisition, project administration, writing-original draft, review and editing. Pieter A.J. Brama: Conceptualization, data curation, funding acquisition, investigation, project administration, supervision, writing-original draft, review and editing.

All authors approved the final version of the manuscript.

## Acknowledgements

The authors acknowledge Yanto Ridwan for the support in the micro-CT scanning, and Mart Verhoeven and Wouter van de Ven for the support in the micro-CT evaluation, and Christian Poinsot for the support in cytotoxicity assessment.

