## Supplement Table 1-4, Figure 1-8 for "Incorporating strontium enriched amorphous calcium phosphate granules in collagen/collagen-magnesium-hydroxyapatite osteochondral scaffold improves subchondral bone repair"

### Supplementary Material

**Table S1:** Clinical Orthopedic Assessment. This tool is to evaluate the clinical health of goat joints.

| Parameter | Variables | Score |
|---|---|---|
| Lameness | Walks normally | 5 |
| | Slightly lame when walking | 4 |
| | Moderately lame when walking | 3 |
| | Severely lame when walking | 2 |
| | Reluctant to rise and will not walk more than five paces | 1 |
| Joint mobility | Full range of motion | 5 |
| | Mild limitation (10–20%) in ROM; no crepitus | 4 |
| | Mild limitation (10–20%) in ROM; with crepitus | 3 |
| | Moderate limitation (20–50%) in ROM; ±crepitus | 2 |
| | Severe limitation (>50%) in ROM; ±crepitus | 1 |
| Pain on knee palpation and movement | None | 5 |
| | Mild signs; Goat turns head in recognition | 4 |
| | Moderate signs; Goat pulls limb away | 3 |
| | Severe signs; Goat vocalises or becomes aggressive | 2 |
| | Goat will not allow palpation | 1 |
| Weight-bearing | Equal on all limbs standing and walking | 5 |
| | Normal standing; favours affected limb when walking | 4 |
| | Partial weight-bearing standing and walking | 3 |
| | Part. weight-bearing standing; non-weight-bearing walk | 2 |
| | Non-weight-bearing standing and walking | 1 |
| Overall score of clinical condition | Not affected | 5 |
| | Mildly affected | 4 |
| | Moderately affected | 3 |
| | Severely affected | 2 |

| | | |
|---|---|---|
| | Very severely affected | 1 |
| Total score | | 25 |

**Table S2:** Macroscopic joint Assessment. This tool is to evaluate the macroscopic normalization of goat joints when the joints were opened.

| **Parameter** | **Variables** | **Score** |
|---|---|---|
| Wound healing abnormal | Yes | 0 |
| | No | 1 |
| Swelling of joints area | Yes | 0 |
| | No | 1 |
| Effusion of the joints | Yes | 0 |
| | No | 1 |
| Patellar luxation | Yes | 0 |
| | No | 1 |
| Joint mobility abnormal / Contractures | Yes | 0 |
| | No | 1 |
| Adhesions of whole joint | Yes | 0 |
| | No | 1 |
| Erosions of whole joint | Yes | 0 |
| | No | 1 |
| Synovial fluid abnormal | Yes | 0 |
| | No | 1 |
| Synovial membrane abnormal | Yes | 0 |
| | No | 1 |
| Lesion on the opposite cartilage surface | Yes | 0 |
| (trochlear ridge vs patella) | No | 1 |
| Lesion on the opposite cartilage surface | Yes | 0 |
| (medial femoral condyle vs meniscus/tibia plateau) | No | 1 |
| | **Total** | **0-11** |

**Table S3:** International Cartilage Repair Society (ICRS) cartilage repair scoring system. This tool is to evaluate the macroscopic appearance of cartilage repair tissue.

| Parameter | Variables | Scores |
|---|---|---|
| Degree of defect repair | In level with surrounding cartilage | 4 |
| | 75% repair of defect depth | 3 |
| | 50% repair of defect depth | 2 |
| | 25% repair of defect depth | 1 |
| | 0% repair of defect depth | 0 |
| Integration to border zone | Complete integration with surrounding cartilage | 4 |
| | Demarcating border < 1 mm | 3 |
| | ¾ of graft integrated, ¼ with a notable border > 1 mm | 2 |
| | 1/2 of graft integrated with surrounding cartilage,1/2 with a notable border > 1 mm | 1 |
| | From no contact to ¼ of graft integrated with surrounding cartilage | 0 |
| Macroscopic appearance | Intact smooth surface | 4 |
| | Fibrillated surface | 3 |
| | Small, scattered fissures or cracks | 2 |
| | Several, small or few but large fissures | 1 |
| | Total degeneration of grafted area | 0 |
| **Overall** | Grade I normal | 12 |
| | Grade II nearly normal | 11-8 |
| | Grade III abnormal | 7-4 |
| | Grade IV severely abnormal | 3-1 |

**Table S4:** A semi-quantitative macroscopic scoring system developed by Goebel et al. for the macroscopic description of articular cartilage repair.

| Parameter | Variables | Scores |
|---|---|---|
| Color of the repair tissue | Hyaline or white | 4 |
| | Predominantly white (>50%) | 3 |
| | Predominantly translucent (>50%) | 2 |
| | Translucent | 1 |
| | No repair tissue | 0 |

| | | |
|---|---|---|
| Presence of blood vessels in the repair tissue | No | 4 |
| | Less than 25% of the repair tissue | 3 |
| | 25-50% of the repair tissue | 2 |
| | 50-75% of the repair tissue | 1 |
| | More than 75% of the repair tissue | 0 |
| Degeneration of adjacent articular cartilage | Normal | 4 |
| | Cracks and/or fibrillations in integration zone | 3 |
| | Diffuse osteoarthritic changes | 2 |
| | Extension of defect into the adjacent cartilage | 1 |
| | Subchondral bone damage | 0 |
| Surface of the repair tissue | Smooth, homogeneous | 4 |
| | Smooth, heterogeneous | 3 |
| | Fibrillated | 2 |
| | Incomplete new repair tissue (rough) | 1 |
| | No repair tissue | 0 |
| Percentage defect filling | 80-100 % | 4 |
| | 60-80 % | 3 |
| | 40-60 % | 2 |
| | 20-40% | 1 |
| | 0-20 % | 0 |
| **Total Scores** | | **20** |

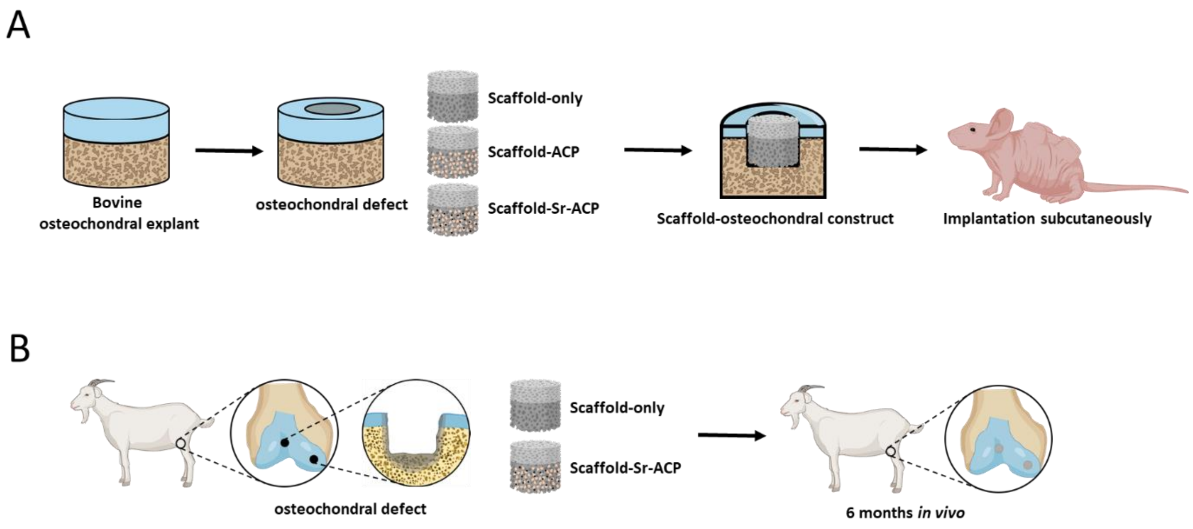


Figure S1. Experiment setup of *in vivo* studies. (A) Scheme of the *in vivo* osteochondral defect mouse model. (C) Scheme of the *in vivo* osteochondral defect goat model.

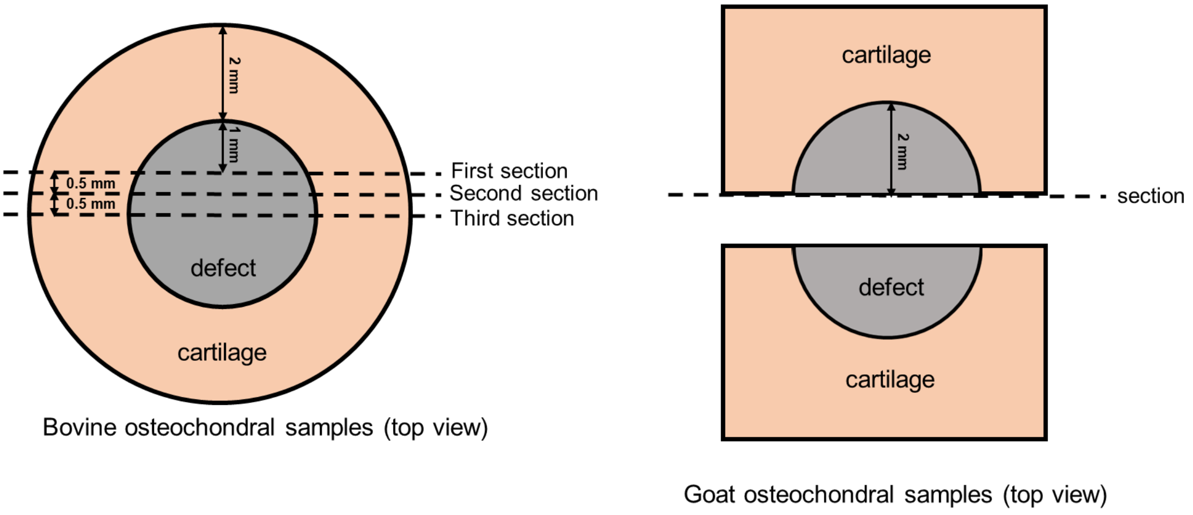


Figure S2. Collection of sections from bovine or goat samples for histology.

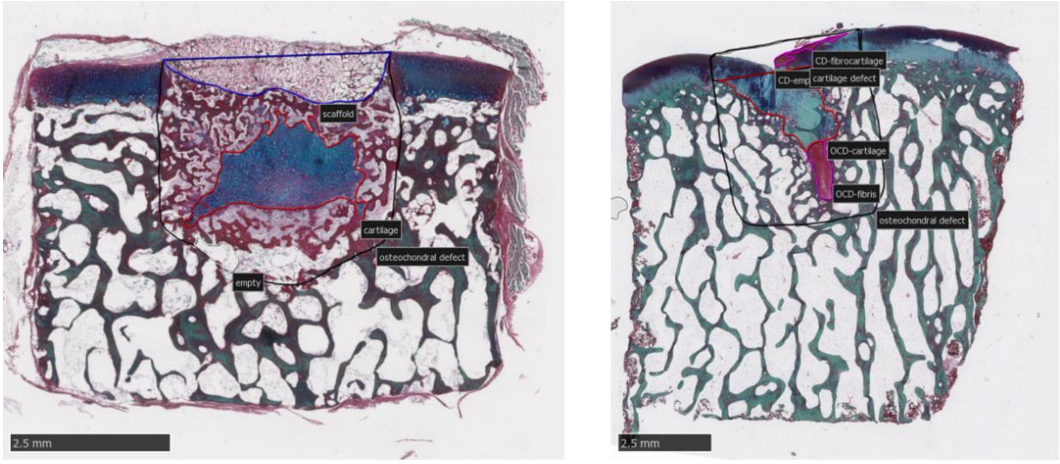


Bovine osteochondral samples　Goat osteochondral samples

Figure S3. Example on defining the defect region, newly formed cartilage-like tissue formation, bone-like tissue formation, fibrous-like tissue formation, remnants of the scaffold for quantification. Scale bars indicated 2.5 mm.

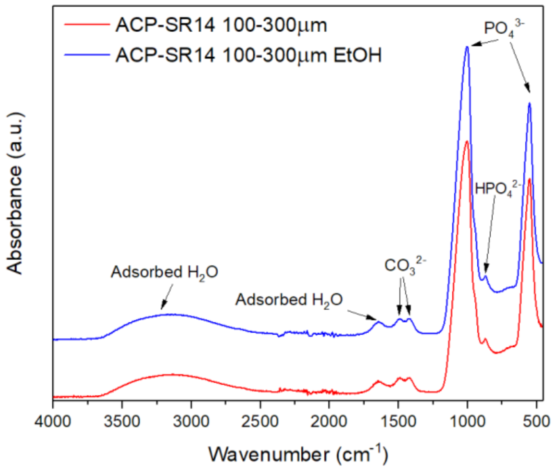


Figure S4: FT-IR spectra of ACP granules before and after rinsing in EtOH to remove debris resulted from dry milling process.

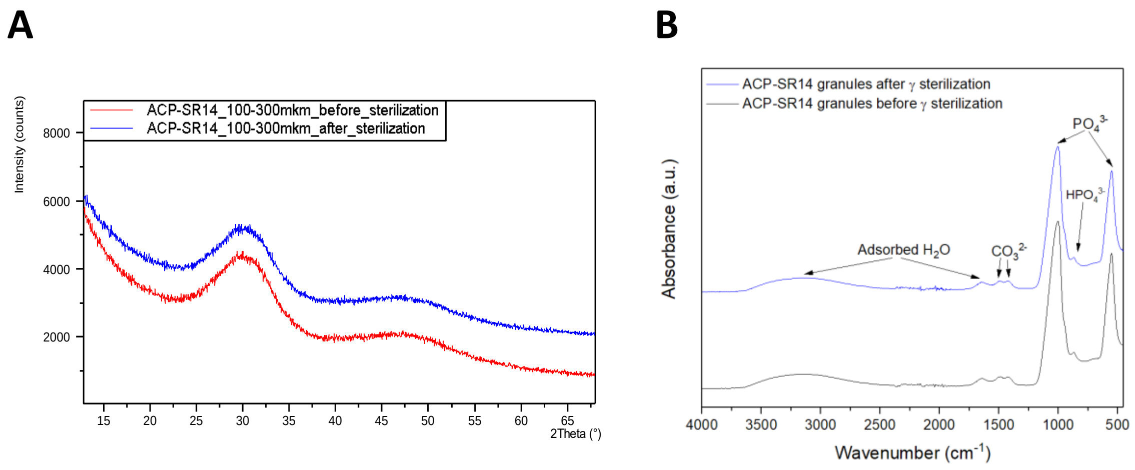


Figure S5: XRD patterns (A) and (FT-IR) spectra of ACP granules before and after γ sterilization.

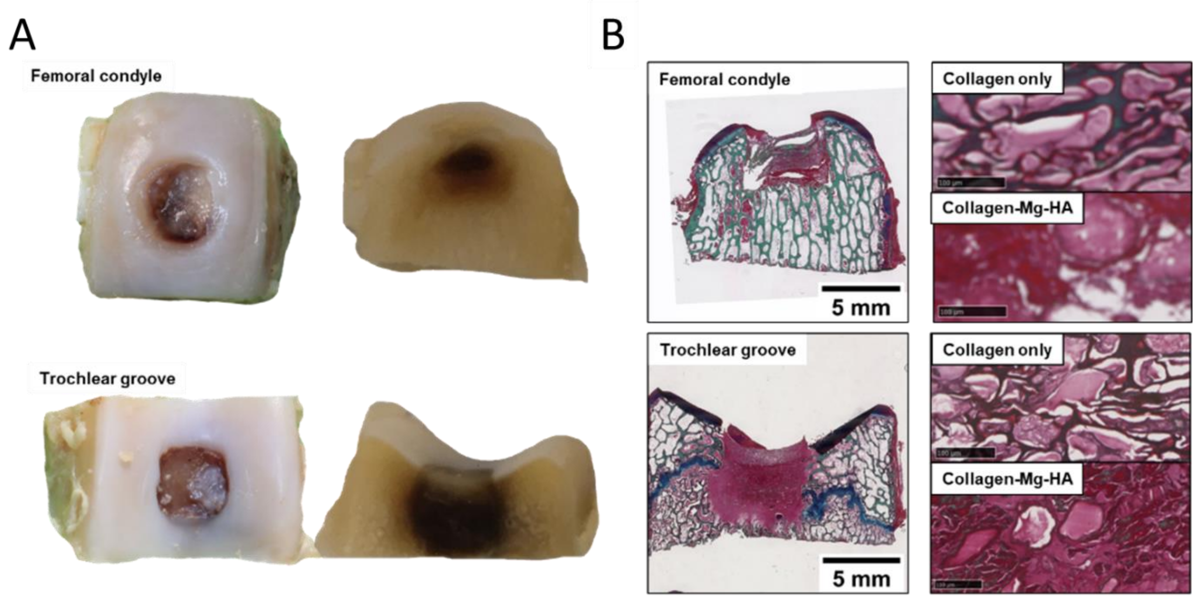


Figure S6: (A) The macroscopic appearance of a femoral condyle defect and a trochlear groove defect 2 weeks after implantation. (B) Two layers of the scaffold implanted in the femoral condyle defect after 2 weeks (stained with Alcian Blue, Fast Green, and Picrosirius Red). The scale bar indicated 5 mm and 100 µm.

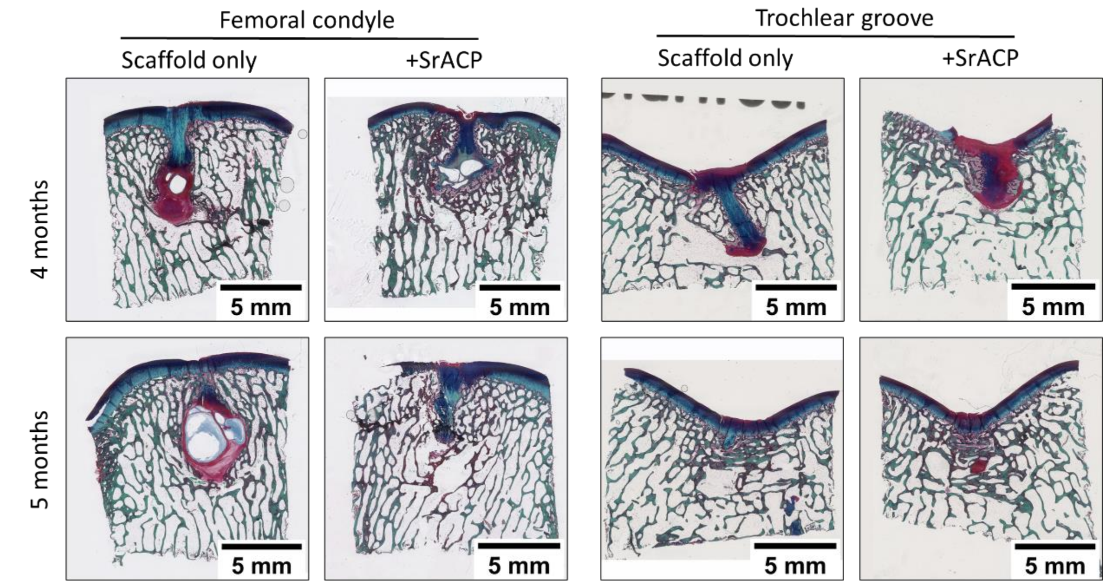


Figure S7: Osteochondral repair at 4- or 5-month post-surgery. RGB (Alcian Blue, Fast Green, and Picrosirius Red) staining of femoral condyle defects and trochlear groove defects treated with either scaffold-only or scaffold + Sr-ACP.

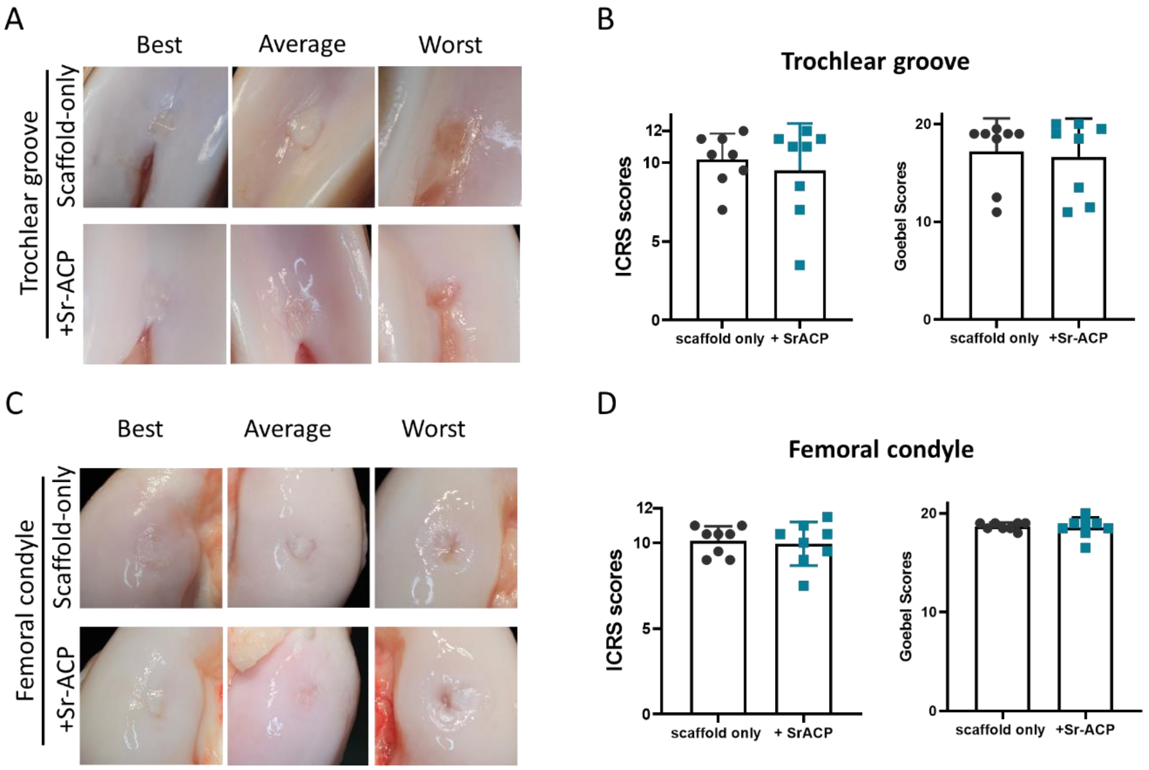


Figure S8: Macroscopic assessment of trochlear groove and femoral condyle defect repair. (A) Representative examples of trochlear groove defect sites treated with scaffold-only or Sr-ACP enriched scaffold after 6 months. Best, average, and worst samples were determined according to the ICRS scores. (B) Macroscopic scores of repair tissue in the trochlear groove defects. (A) Representative examples of femoral condyle defect sites treated with scaffold only or scaffold with Sr-ACP after 6 months. Best, average, and worst samples were determined according to the ICRS scores. (C) Macroscopic scores of repair tissue in femoral condyle defects. The maximum score for ICRS is 12 (indicating the best), and the maximum score for Goebel score is 20 (indicating the best).
